# Quantification of gene expression patterns to reveal the origins of abnormal morphogenesis

**DOI:** 10.1101/246256

**Authors:** N. Martínez-Abadías, R. Mateu, J. Sastre, S Motch Perrine, M Yoon, A. Robert-Moreno, J. Swoger, L. Russo, K Kawasaki, J. Richtsmeier, J. Sharpe

**Affiliations:** Centre for Genomic Regulation (CRG), The Barcelona Institute for Science and Technology, Dr. Aiguader 88, 08003 Barcelona, Spain; Universitat Pompeu Fabra (UPF), Barcelona, Spain; Universitat de Barcelona, Barcelona, Spain; Universitat de les Illes Balears (UIB), Palma de Mallorca, Spain; Pennsylvania State University, University Park (PA), USA; Institució Catalana de Recerca i Estudis Avançats (ICREA), Barcelona, Spain

**Keywords:** Developmental defects, Optical Projection Tomography (OPT), Whole-Mount-in-Situ Hybridization (WISH), Geometric Morphometrics (GM), quantitative shape analysis, limb development, Apert syndrome

## Abstract

The earliest developmental origins of dysmorphologies are poorly understood in many congenital diseases. They often remain elusive because the first signs of genetic misregulation may initiate as subtle changes in gene expression, which can be obscured later in development due to secondary phenotypic effects. We here develop a method to trace back the origins of phenotypic abnormalities by accurately quantifying the 3D spatial distribution of gene expression domains in developing organs. By applying geometric morphometrics to 3D gene expression data obtained by Optical Projection Tomography, our approach is sensitive enough to find regulatory abnormalities never previously detected. We identified subtle but significant differences in gene expression of a downstream target of the *Fgfr2* mutation associated with Apert syndrome. Challenging previous reports, we demonstrate that Apert syndrome mouse models can further our understanding of limb defects in the human condition. Our method can be applied to other organ systems and models to investigate the etiology of malformations.

---

Morphogenesis is guided by dynamic spatio-temporal regulation of gene expression patterns^1,2^, the regions within tissues where genes are expressed at specific developmental times. Critical changes in the space, time or intensity of gene expression patterns can result in organ malformation, with reduced or even loss of function. These errors of morphogenesis, which occur in approximately 3% of live births^3^, are induced by environmental and/or genetic insults that alter the normal process of development. The common approach to reveal the origin of these alterations is to look for phenotypic abnormalities in animal models and visually assess the overall patterns of gene expression. However, this qualitative approach has not been able to identify the earliest signs of dysmorphogenesis in many diseases because the first changes in the gene expression patterns may be subtle, limited in time, and later abnormalities may obscure the original genetic cause. To reveal the primary etiology of congenital malformations, more rigorous methods are needed to quantify the 3D phenotypes of gene expression patterns. A useful tool to trace back development should be able to perform a quantitative statistical comparison of normal and disease-altered embryogenesis and detect the earliest signs of genetic misregulation leading to organ malformation.

Developing organs are shaped by regulating gene activity in space and time^1^. Cascades of gene regulatory networks provide the detailed instructions to organize cell behaviour and orchestrate tissue growth and differentiation. Gene expression patterns can be readily mapped within tissues in a true three-dimensional framework combining Whole-Mount-in-Situ Hybridization (WISH)^4^ with Optical Projection Tomography (OPT)^5^. Whereas WISH is a standard molecular technique for detecting the expression of a specific gene using a labelled complementary RNA probe^6,7^, OPT is a mesoscopic imaging procedure that can produce high resolution 3D reconstructions of whole developing embryos processed by WISH^8,9^. These technologies represented breakthroughs in developmental biology and have provided invaluable qualitative insights into gene function and development^8^. However, methods for quantifying the 3D gene expression distributions in a systematic, objective manner are still lacking.

Expanding the potential of OPT from *qualitative* to *quantitative* analysis of gene expression patterns is challenging. The expression of genes is characterized by highly dynamic patterns, with fast rates of change over time and fuzzy boundaries that usually do not correspond to well-defined anatomical structures but to tissue regions where cells are dynamically up- and down-regulating genes. The quantification of gene expression patterns has been rarely done^10–16^. We propose to quantify the shape of developing organs in association with their underlying gene expression patterns applying Geometric Morphometrics (GM), a set of statistical tools for measuring and comparing shapes with increased precision and efficiency^17–22^. This approach provides the ability to replace qualitative observations with quantification of subtle yet significant biological differences that underlie the processes of morphogenesis altered by disease.

Here we illustrate how our method can reveal the genetic origin of developmental defects by investigating limb malformations in Apert syndrome [OMIM 101200]. Apert syndrome is a rare congenital disease, with disease prevalence of 15–16 per million live births, that is characterized by cranial, neural, limb, and visceral malformations^23^. Over 99% of Apert cases are associated with one of two missense mutations, S252W and P253R, on Fibroblast Growth Factor Receptor 2 (FGFR2)^24,25^. The mutations occur on neighbouring amino acids on the linker region between the second and third extracellular immunoglobulin domains of FGFR2, and alter the ligand-binding specificity of the receptors^26,27^. Thus, the FGF receptors are activated inappropriately, altering the entire FGF/FGFR signalling pathway and causing dysmorphologies of different organs and systems^28^. Apert syndrome shares craniofacial dysmorphologies with other craniosynostosis syndromes but is differentiated on the basis of limb defects of fore- and hindlimb digits. The craniofacial dysmorphology of Apert syndrome (i.e. premature closure of cranial sutures and patent anterior fontanelle associated with atypical head shape, midfacial retrusion and palatal defects)^23^ has been intensively investigated, especially because mouse models show cranial phenotypes that correspond with the human condition^29–33^. However, the associated limb defects are less well studied, in part because mouse models for Apert syndrome present only subtle limb anomalies^34–36^, even in those carrying the FGFR2 P253R mutation^34^ that is associated with the more severe limb malformations in Apert syndrome^37,38^.

Here we present precise phenotyping of limbs of newborn and embryonic specimens of the *Fgfr2*^*+/P253R*^ Apert syndrome mouse model and reveal significant differences that can be traced to as early as one day after the initiation of limb development. To explore the molecular basis of these initial signs of limb dysmorphology we applied our method combining OPT and GM to assess the expression pattern of a direct target of *Fgf* signalling, *Dusp6*, a relevant gene for limb morphogenesis^39,40^. Our quantitative analyses demonstrate that the Apert syndrome *Fgfr2 P253R* mutation induces changes in the expression pattern of *Dusp6* and that these genetic changes are associated with significant phenotypic alterations. These results provide insight into the origins of limb malformations in Apert syndrome.

## Results

### Apert syndrome mice present limb malformations at birth

Previous studies reported that Apert syndrome mice do not show obvious abnormalities of the limb^35^, and thus focused their molecular analyses on the skull^34^. Histopathological analyses in Apert syndrome mice only revealed overall limb shortening due to abnormal osteogenic differentiation, but no signs of limb disproportion or syndactyly^34,36^. As a further test of whether or not the *Fgfr2 P253R* mutation affects limb development in mice, we first performed an extensive quantitative analysis of the size and shape of individual forelimb bones using data from high resolution microCT images of newborn (P0) mutant and unaffected littermates (Fig. 1a-f). Our results revealed many more significant differences between P0 unaffected and *Fgfr2*^*+/P253R*^ mutant littermates than previously reported. We found that the humerus, radius and ulna were statistically significantly shorter in length but had increased bone volumes in *Fgfr2*^*+/P253R*^ mutant mice in comparison to unaffected littermates (Fig. 1g and Table SI_1). More localized size differences were detected in the bones derived from the autopod that give rise to the hands. The distal phalanx of digit I, the proximal phalanx of digit V, and metacarpals II, III and IV were significantly longer in *Fgfr2*^*+/P253R*^ mutant mice (Fig. 1g and Table SI_1). In contrast, the proximal phalanx of digit III was shorter and lower in bone volume in *Fgfr2*^*+/P253R*^ Apert syndrome mice relative to unaffected littermates (Fig. 1g and Table SI_1). The scapula and the clavicle, the bones that form the shoulder girdle, were also significantly affected: the scapula was longer, the clavicle was shorter and both bones showed increased bone volumes in *Fgfr2*^*+/P253R*^ Apert syndrome mice (Fig. 1g and Table SI_1) compared to unaffected littermates.

**Figure 1.**
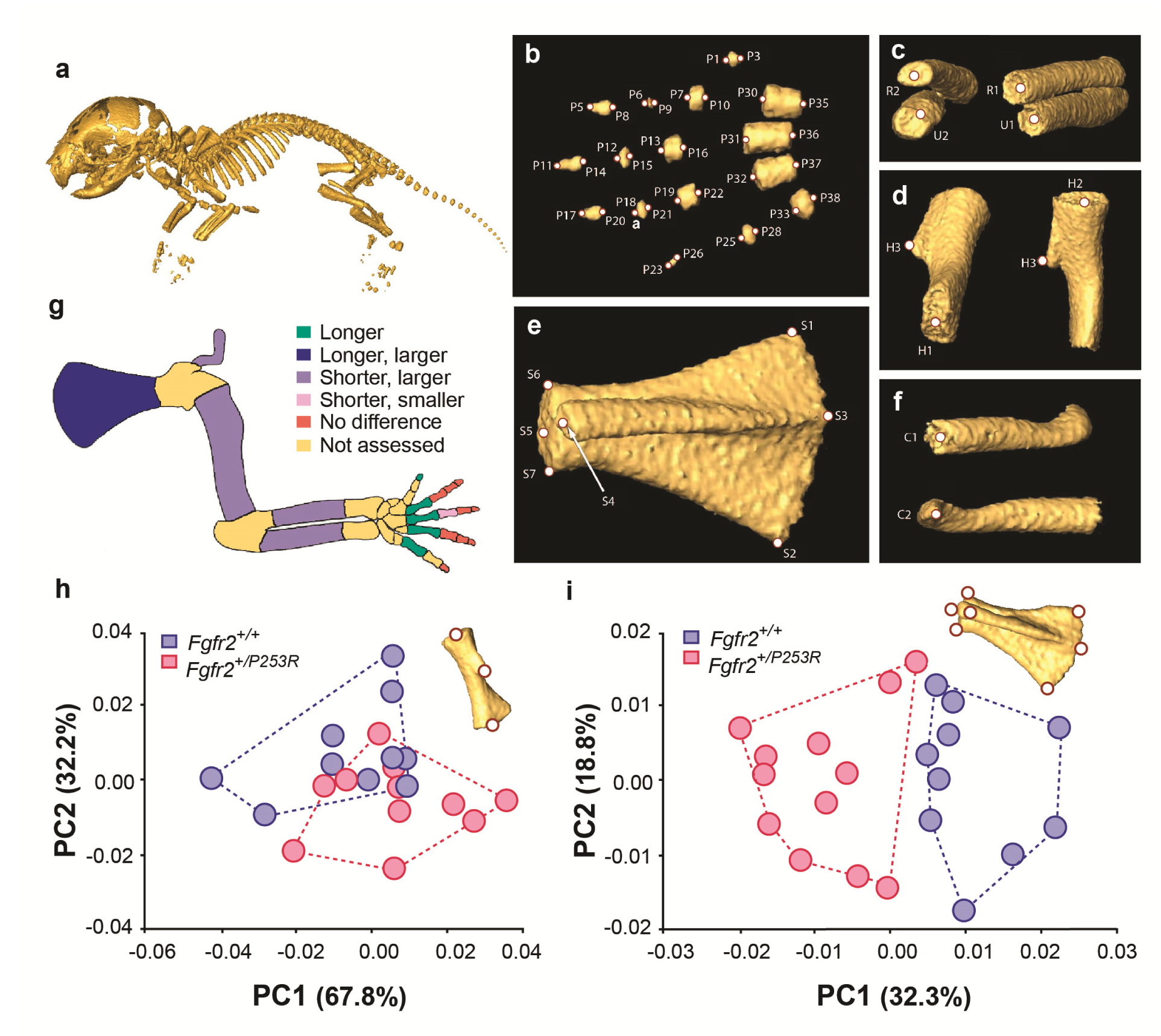
Quantitative size and shape comparison of forelimb bones in *Fgfr2*^*+/P253R*^ newborn mice (P0) and unaffected littermates (P0). a) Mouse skeleton at P0. 3D isosurface reconstruction of the skeleton of an unaffected littermate obtained from a high resolution µCT scan. b-f) Anatomical landmarks recorded on µCT scans of Apert syndrome mice at P0. b) Autopod (hand). Landmarks were recorded at the midpoint of the proximal and distal tips of the distal, mid and proximal phalanges (P1-P28) and the metacarpals (P29-P38). Proximal phalanx I, middle phalanx V, and metacarpal I are not displayed because these bones have not yet mineralized at P0 and could not be visualized in many specimens. c) Zeugopod. Landmarks were recorded at the midpoint of the proximal and distal tips of the radius (R1- R2) and ulna (U1-U2). d) Stylopod. Landmarks were recorded at the midpoint of the proximal and distal tips of the humerus (H1-H2), as well as the tip of the deltoid process (H3). e) Scapula. Landmarks were recorded at the most superior and inferior lateral points of the scapula (S1-S2), the most posterior point of the spine (S3), most antero-medial point of the acromion process (S4) and the medial, superior and inferior points of the glenoid cavity (S5-S7). f) Clavicle. Landmarks were recorded at the medial point of the sternal and the acromial ends (C1-C2). g) Length and volume differences in the forelimbs of Apert syndrome mouse models. Schematic representation of the forelimb of a P0 mouse showing in different colours statistically significant differences in bone length and volume as measured by two-tailed one-way ANOVA or Mann-Whitney U-test in *Fgfr2*^*+/P253R*^ mice relative to unaffected littermates as specified in Table SI_1. Longer/shorter refer to length, whereas larger/smaller refer to volume. Blue: Longer and larger, green: longer, purple: shorter and larger, pink: shorter and smaller, orange: no difference, yellow: not assessed because bones were not visible/ossified. h, i) Shape differences in the forelimbs of Apert syndrome mouse models. Scatterplots of PC1 and PC2 scores based on Procrustes analysis of anatomical landmark locations representing the shape of the left humerus (h) and the left scapula (i) of unaffected (n=10) and mutant (n=12) littermates of Apert syndrome mouse models. Convex hulls represent the ranges of variation within each group of mice.

The PCA based on the shape of the humerus did not show marked shape differences between unaffected and *Fgfr2*^*+/P253R*^ Apert syndrome mice (Fig. 1h). However, the PCA of the scapula indicated a clear morphological differentiation between groups (Fig. 1i). The scapula of *Fgfr2*^*+/P253R*^ Apert syndrome mice presented a more robust phenotype, with wider and longer scapulae in comparison to their unaffected littermates.

Overall, these size and shape differences demonstrate that *Fgfr2*^*+/P253R*^ Apert syndrome mice present widespread and significant limb dysmorphologies at P0 that were not previously reported and would not have been revealed without quantitative statistical testing. Some defects have a direct correspondence with the human phenotype, such as shoulder anomalies and short humeri^24^. However, in newborn mice we did not detect any clear sign of syndactyly, which is the most prominent limb defect in people with Apert syndrome^41,42^. Since the forelimb of mice is not yet completely ossified at P0 and *Fgfr2*^*+/P253R*^ mutant littermates die shortly after birth, we could not assess whether other limb abnormalities appear later in development.

### A quantitative morphometric method to assess embryonic gene expression patterns

To determine the developmental basis of the limb anomalies quantified in newborn mice, we developed a quantitative method to explore early embryonic limb development. First, to visualize the expression pattern of a downstream target of *Fgfr2*, we obtained OPT scans of *Fgfr2*^*+/P253R*^ Apert syndrome mouse embryos analysed with WISH to reveal *Dusp6* expression (Fig. 2). Qualitative assessment of the 3D reconstructions showed that *Dusp6* was widely expressed throughout the embryo from embryonic day (E) 10.5 to E11.5, with highest intensity in the limbs, the head and the spinal cord (Fig. 2). *Dusp6* was also expressed in the heart with moderate intensity. Visually comparing the distribution of the *Dusp6* gene expression pattern it was possible to distinguish between embryos at E10.5 and E11.5 stages of development. At E10.5, *Dusp6* was prominently expressed in the facial prominences and along the entire spinal cord, whereas at E11.5 the expression of *Dusp6* was more widespread in the brain and limited to the tail. Focusing on the limbs, the expression of *Dusp6* at the two different stages was also readily distinguishable, with *Dusp6* expression domains thinning into a more extended domain along the limb outline as the limb buds grow from E10.5 to E11.5 (Fig. 2). However, due to large amount of developmental variation within litters, *Fgfr2*^*+/P253R*^ mutant and unaffected littermates were not distinguishable from each other (Fig. 2).

**Figure 2.**
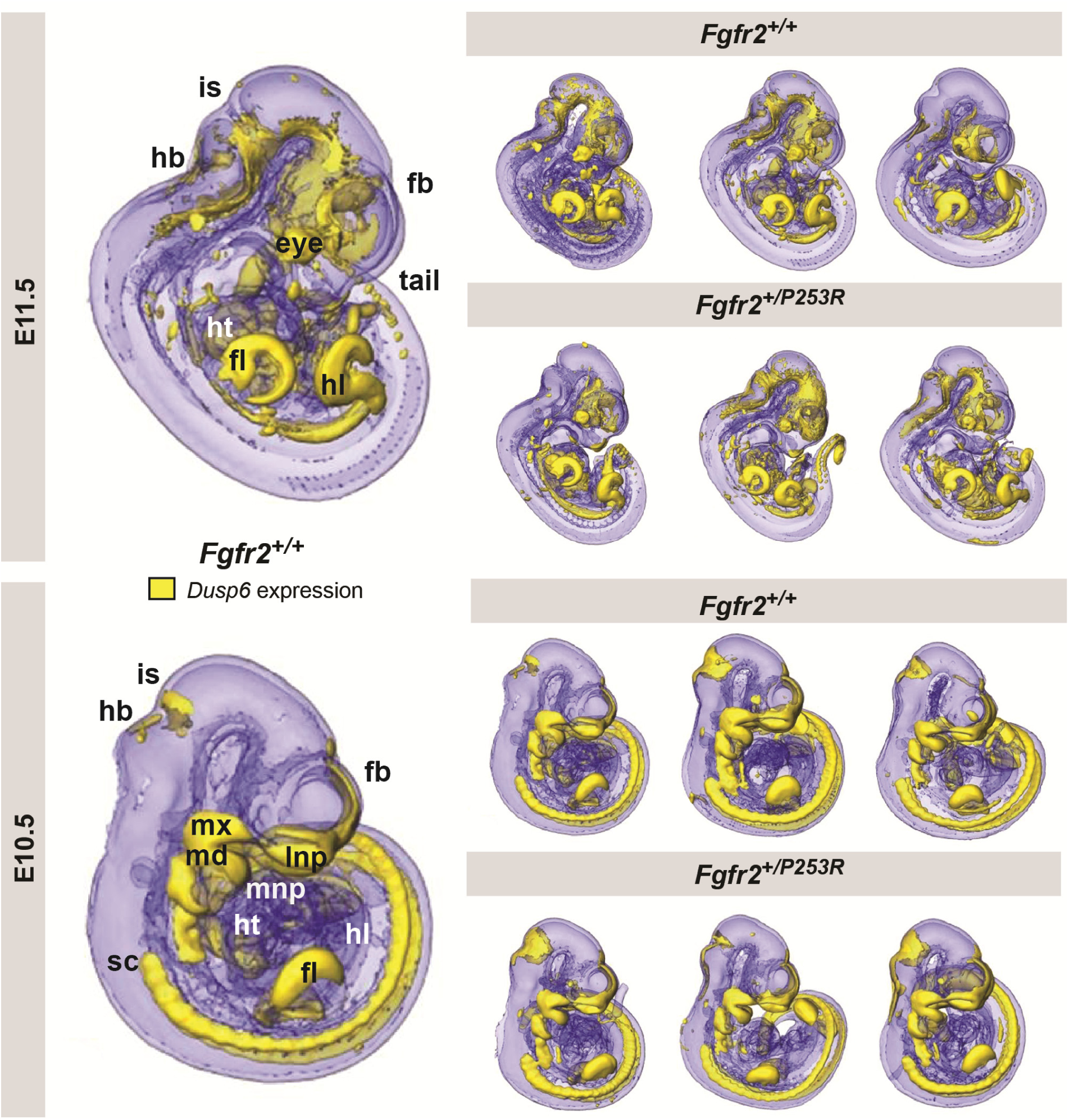
Qualitative visualisation of gene expression of *Dusp6* in unaffected and *Fgfr2*^*+/P253R*^ mouse embryos at E10.5 and E11.5. OPT scans of embryos WISH stained for *Dusp6* revealed the anatomical location of gene expression (shown in yellow). For each stage, the main expression domains are highlighted on the left for anatomical reference on a lateral view of a 3D reconstruction of a *Fgfr2*^*+/+*^ unaffected embryo (fb: forebrain, is: isthmus, hb: hindbrain, mx: maxillary prominence, md: mandibular prominence, lnp: lateral nasal process, mnp: medial nasal process, ht: heart, sc: spinal cord, fl: forelimb, hl: hindlimb). On the right, 3D reconstructions of three unaffected and three *Fgfr2*^*+/P253R*^ mutant embryos from the same litter are displayed to represent the high degree of variation in developmental age within litters. Embryos are not to scale.

Quantitative testing was thus required to more accurately evaluate limb alterations potentially associated with Apert syndrome but undetectable by eye. We developed a method for 3D shape analysis of the limb and associated gene expression pattern of *Dusp6* (Fig. 3 and Video SI_1). This protocol enabled us to determine differences in limb size and shape between genotype groups and whether these phenotypic differences are associated with altered gene expression patterns (Fig. 3). Our approach uses GM methods to directly measure the limb anatomy and gene expression domains segmented from the 3D reconstructions of the embryo OPT scans. As expression of *Dusp6* showed a fuzzy spatial gradient, multiple thresholding was used to consistently define a high gene expression pattern (Fig. 3, steps from 1 to 5). After manual and semiautomatic recording of 3D coordinates of landmarks on the surfaces of the limb and the gene expression domains blinded to group allocation (Fig. 3, step 6), multivariate statistical analyses were performed to explore shape and size variation and covariation patterns between the limb morphology and the *Dusp6* domain (Fig. 3, steps 7 and 8).

**Figure 3.**
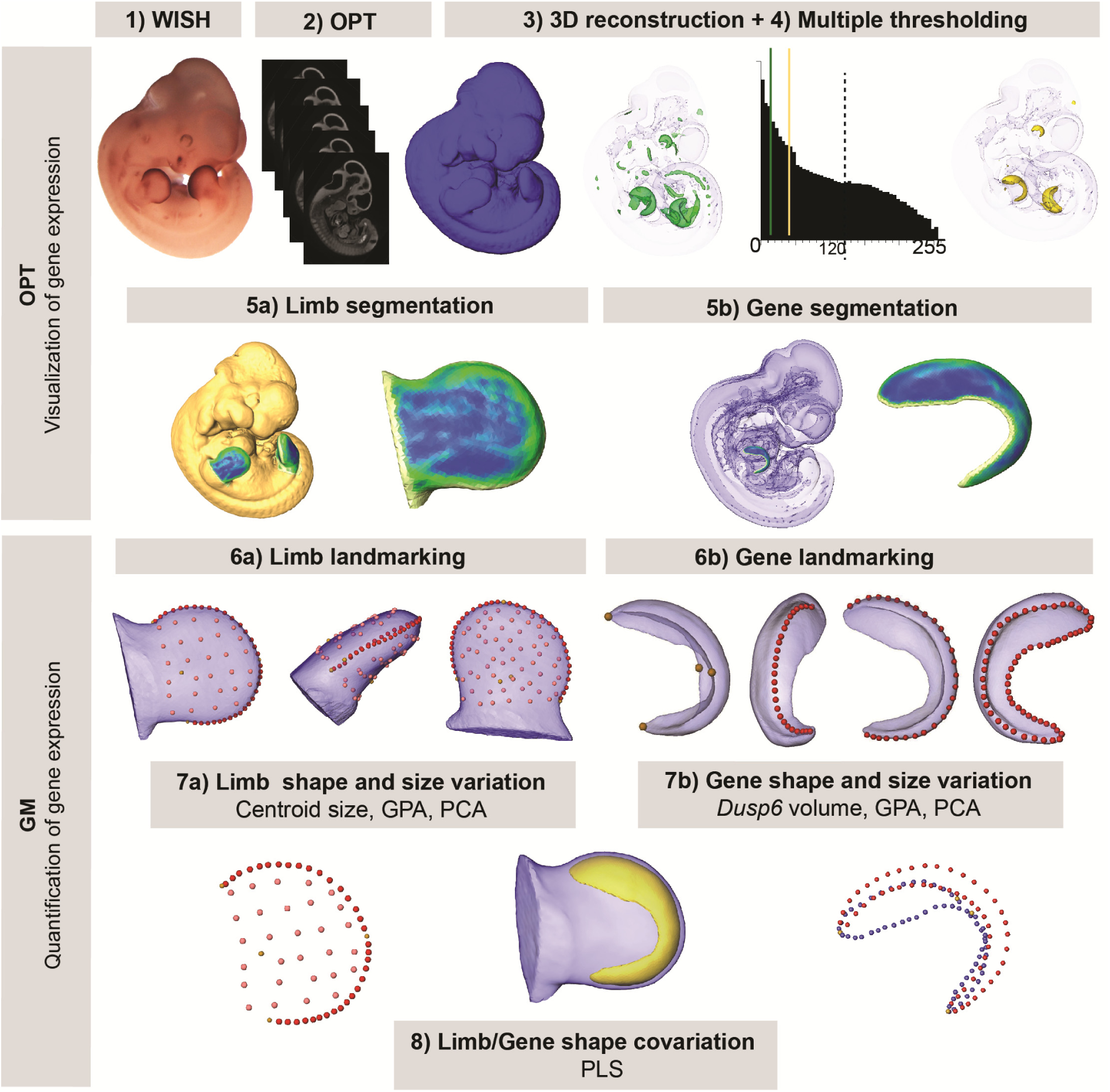
New quantitative analysis method for 3D gene expression data, based on geometric morphometrics. Mouse embryos between E10.5 and E11.5 were analysed with WISH to reveal the expression of *Dusp6* (1), and then cleared with BABB and OPT scanned using both fluorescence and transmission light (2). The external surface of the embryo was obtained from the 3D reconstruction of the fluorescence scan (2). Multiple thresholding of the transmission scan by choosing different levels of grey values as shown by the histogram allowed to visualize gene expression patterns at different intensities (3). Moderate gene expression (shown in green) was displayed as the isosurface obtained using as a threshold the grey value computed as 2/3 of the last grey value showing the *Dusp6* expression domain (3). High gene expression (shown in yellow) was displayed as the isosurface obtained using as a threshold the grey value computed as 1/3 of the last grey value showing the *Dusp6* expression domain (3). From the whole mouse embryo isosurfaces, all four limb buds were segmented (5a). From the high gene expression isosurface, the *Dusp6* domains from all the available limbs were segmented (5b). Maximum curvature patterns were displayed to optimize landmark recording (5). For each limb we captured the shape and size of the limb bud (6a) and the underlying high *Dusp6* gene expression pattern (6b) recording the 3D coordinates of anatomical landmarks (yellow dots), curve semilandmarks (red dots) and surface semilandmarks (pink dots). Anatomical and curve landmarks were recorded manually on each limb. Surface landmarks were recorded on one template limb and interpolated onto target limbs. Landmark coordinates were the input for Geometric Morphometric (GM) quantitative shape analysis (7, 8) to superimpose the landmark data (GPA, General Procrustes Analysis), compute limb size (Centroid size), and explore shape variation within limbs and gene expression domains by litters (PCA, Principal Component Analysis). Finally, the covariation patterns between the shape of the limb and the shape of the gene expression domain were also explored (PLS, Partial Least squares).

### The first signs of limb dysmorphology in Apert syndrome

Since gene expression patterns are highly dynamic and rapidly change in size, shape and position within a few hours of development^15^, individual limb buds from *Fgfr2*^*+/P253R*^ mutant embryos and their unaffected littermates aged between E10.5 and E11.5 were staged using a fine-resolution staging system (http://limbstaging.crg.es)^43^. The staging results showed that the analysed limbs represent a temporal continuum over development, with no significant differences between the staging of unaffected and mutant littermates of the same litter (Fig. SI_1). We partitioned the time span from E10 to E11.5 into four periods, each one approximately representing 12 hours of development (see Table SI_2 and Methods for further details on sample composition). The analysis of the complete dataset that considers hindlimbs and forelimbs from each litter separately is available as Supplementary Information.

We first focused on analysing limb dysmorphology, aiming to determine the youngest stage which showed morphological differences between mutant and unaffected limbs. To trace limb development back in time we first analysed embryos from the oldest period (as the differences would be easier to find) and from there proceeded towards the earlier (younger) periods. In this way we should confidently identify the initiation of limb dysmorphogenesis associated with the *Fgfr2 P253R* mutation determining using geometric morphometric methods.

During the “Late” period, we detected that *Fgfr2*^*+/P253R*^ mice were already clearly separated from their unaffected littermates in the morphospace defined by the Principal Component Analysis (PCA) (Fig. 4a). Relative to their unaffected littermates, limbs of *Fgfr2*^*+/P253R*^ mice presented subtle phenotypic limb differences: limbs were shorter, wider and more robust, with limited development of the wrist (Fig. 4a). Quantitative comparison of limb size showed that the limbs of mutant mice were also significantly smaller (Fig. 5a and Table SI_3). Overall, these results confirmed that the *Fgfr2 P253R* Apert syndrome mutation has an effect on limb development, altering both the size and shape of the limbs. Most likely, these subtle but significant phenotypic differences would have remained undetected by a qualitative approach. Our quantitative approach revealed their statistical significance and pointed to the origin of the Apert syndrome limb malformation prior to E11.5, before the “Late” period.

**Figure 4.**
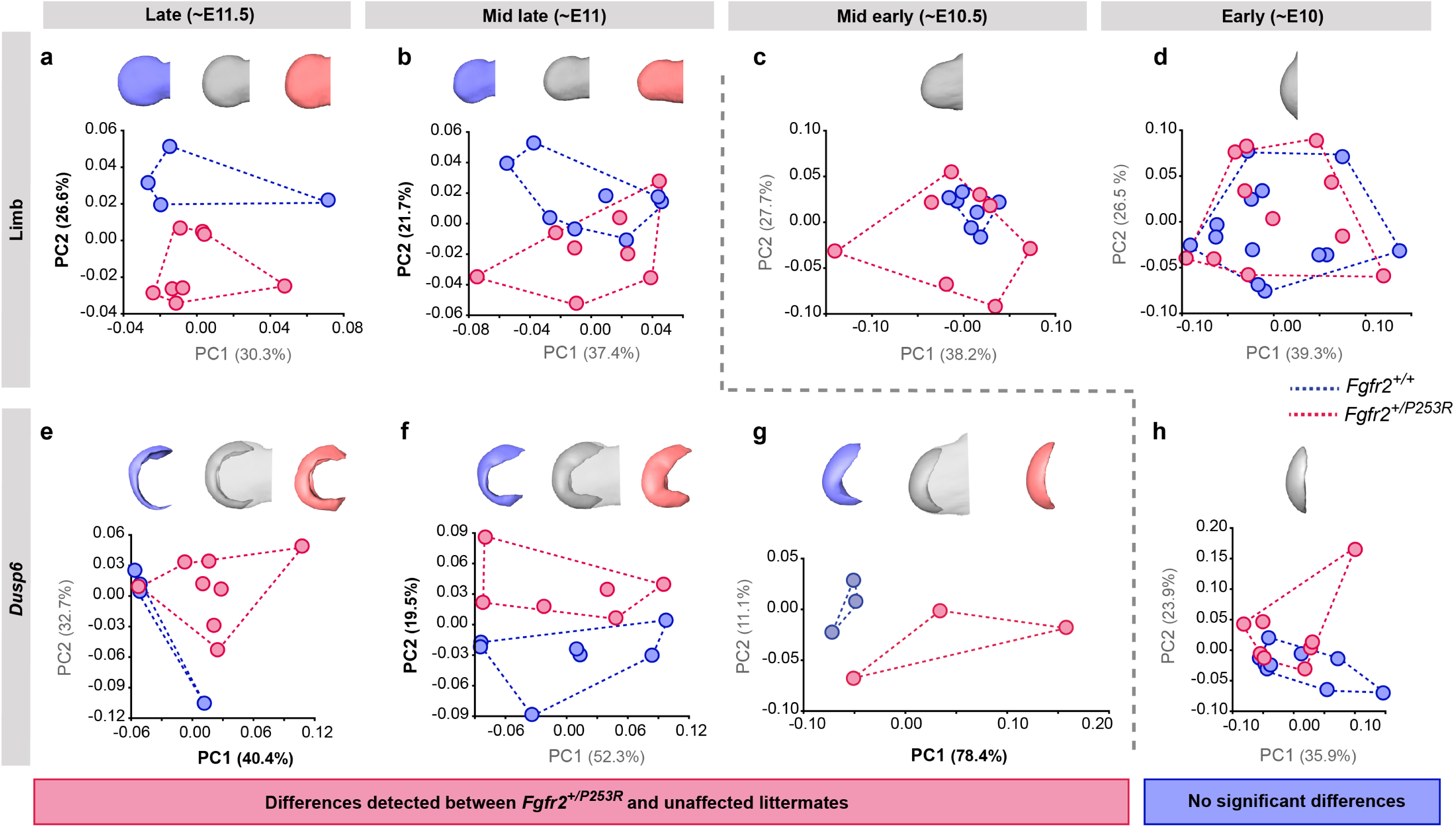
Tracing of limb phenotypes (anatomical and molecular) back through developmental time to the earliest moment of appearance. Principal Component Analyses based on the Procrustes-based semilandmark was used to analyse the shape of the limbs and the corresponding *Dusp6* expression domains at each developmental period. Each period was analysed separately for the shape of the limb (a-d) and the *Dusp6* expression domain (e-h), as specified in Methods and Table SI_2. Scatterplots of PC1 and PC2 axes with the corresponding percentage of total morphological variation explained are displayed for each analysis, along with morphings associated with the negative, mid and positive values of the PC axis that separates mutant and unaffected littermates (PC1 or PC2, as highlighted in bold black letters in the corresponding axis). Morphings are displayed in grey tones when the analysis showed no differentiation between mutant and unaffected littermates. Morphs are displayed in colour when the analysis revealed differentiation between mice (blue: unaffected littermates; pink: mutant littermates). Limb buds and *Dusp6* domains are not to scale and oriented with the distal aspect to the left, the proximal aspect to the right, the anterior aspect at the top and the posterior aspect at the bottom of all images. Convex hulls represent the ranges of variation within each group of mice.

**Figure 5.**
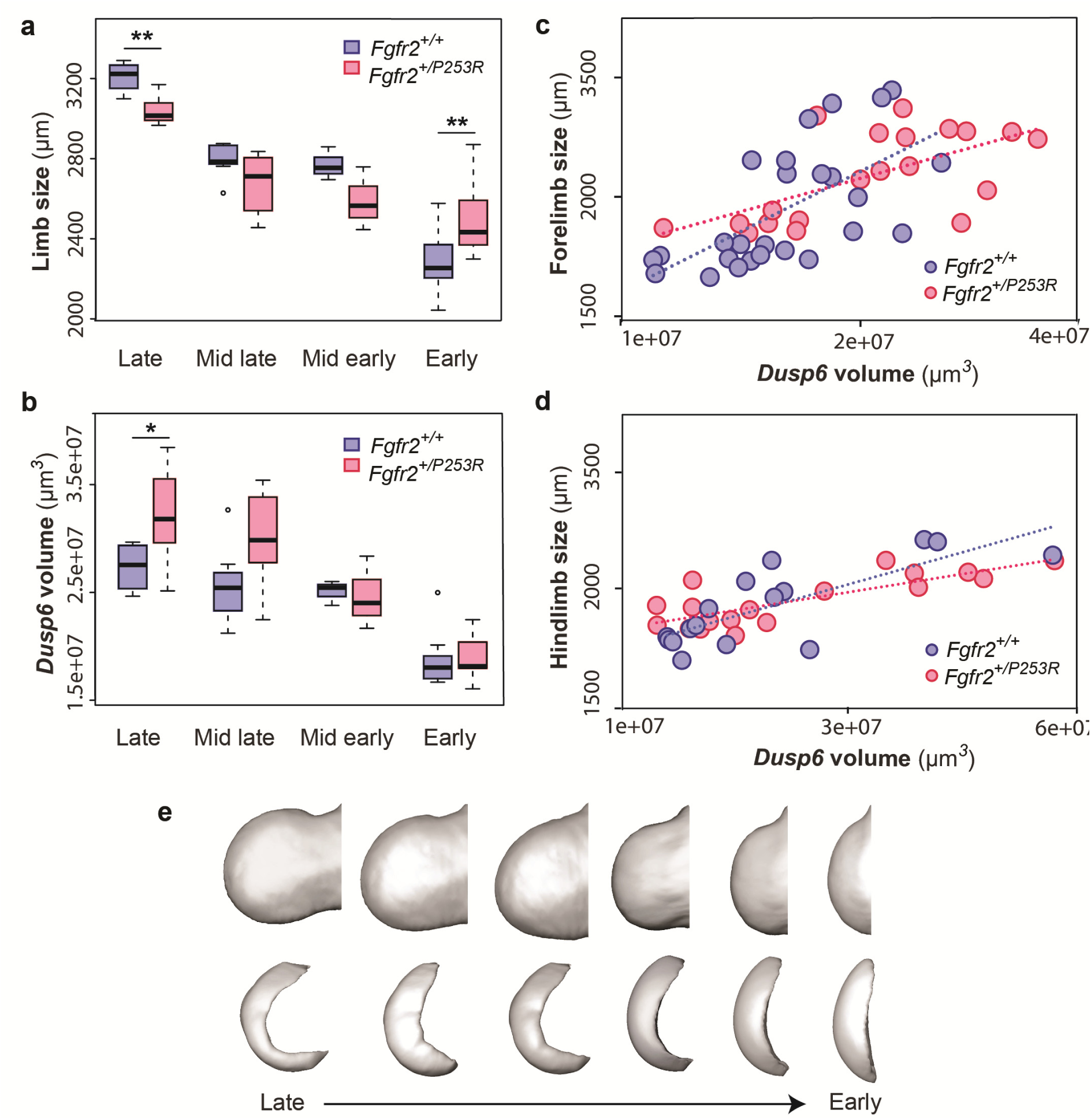
Quantitative correspondence between the size and shape of the limbs and the *Dusp6* expression pattern. a-d) Comparison of limb bud size and Dusp6 volume in unaffected and *Fgfr2*^*+/P253R*^ mutant littermates across development. Limb size was measured as limb centroid size (a), whereas the size of the *Dusp6* expression was measured as the volume of the gene domain (b), as specified in Methods and Tables SI_2 and 3. The association between the size of the limbs and the volume of the *Dusp6* domain was assessed separately for forelimbs (c) and hindlimbs (d). Statistical significant differences as revealed by two-tailed t-tests are marked with asterisks, representing the degree of significance: \**P-value*=0.03, ** *P-value*=0.01. e) Time course assessing the morphological integration pattern between the limb phenotype and the shape of the gene expression pattern using Partial Least Squares analyses. Associated shape changes from late to early limb development are shown from morphings associated with the negative, mid and positive values of PLS1, which accounted for almost all the covariation (97.6% in forelimbs and 99.5% in hindlimbs) between the limb buds and the *Dusp6* gene expression domains. Limb buds and *Dusp6* domains are not to scale and oriented distally to the left, proximally to the right, anteriorly to the top and posteriorly to the bottom. On the right, representing the positive extreme of PLS1 axis, typical early limb buds showed a protruding shape (i.e. short in the proximo-distal axis and symmetrical in the antero-posterior axis) associated with a flat-bean shaped *Dusp6* expression pattern localized in the distal limb region, underlying the apical ectodermal ridge, and spreading proximally towards the dorsal and ventral sides of the limb. On the left, representing the negative extreme of the PLS1 axis, limb buds were elongated in the proximal axis, asymmetric on the antero-posterior axis, with an expansion of the distal limb region and a contraction of the proximal region, at the wrist level. This typical limb shape of more developed limbs was associated with a *Dusp6* expression that was extended underneath the apical ectodermal ridge towards the anterior and the posterior ends of the gene expression, but reduced on the dorsal and ventral sides of the limb.

At the “Mid late” period, the limbs of *Fgfr2*^*+/P253R*^ mutant mice were still distinguishable from the limbs of unaffected mice in the morphospace of the PCA (Fig. 4b). At this period, the limbs of *Fgfr2*^*+/P253R*^ mice lacked the antero-posterior asymmetry and the narrowing of the wrist region more typical of unaffected littermates. Instead, *Fgfr2*^*+/P253R*^ mice showed a limb phenotype that was elongated in the proximo-distal axis and thickened in the dorso-ventral axis (Fig. 4b), resembling the limb shape of younger unaffected embryos. This shape difference coincided with reduced growth in *Fgfr2*^*+/P253R*^ mice, as the limbs of *Fgfr2*^*+/P253R*^ mice tended to be smaller than unaffected limbs (Fig. 5a and Table SI_3). Therefore, significant differences between unaffected and mutant limbs still could be detected at the “Mid late” period of development and the origins of limb defects associated with Apert syndrome should be sought earlier in development.

The first period where no significant differences could be detected between unaffected and *Fgfr2*^*+/P253R*^ Apert syndrome mice was at the “Mid early” period (Fig. 4c). Despite internal variation in limb shape, with *Fgfr2*^*+/P253R*^ mice spreading throughout the morphospace and unaffected littermates concentrated on one region, mutant and unaffected littermates completely overlapped. Therefore, limb shape differences could no longer be detected between groups. Limb size differences were not significant either (Fig. 5a and Table SI_3). Therefore, our results suggest that the critical time point of limb dysmorphogenesis associated with Apert syndrome occurred between the “Mid late” and “Mid early” periods, corresponding to the transition period from E10.5 to E11 (Fig. 4b-c).

Finally, no further sign of limb shape dysmorphology was detected at the “Early” period of development (Fig. 4d). At this period there was a great range of developmental variation, with unaffected and *Fgfr2*^*+/P253R*^ mutant mice completely overlapping in the morphospace and all limbs displaying similar incipient bud shapes (Fig. 4d). The limbs of *Fgfr2*^*+/P253R*^ mice were significantly larger than the limbs of their unaffected littermates (Fig. 5a and Table SI_3), suggesting that at this early time point there is a significant effect of the *Fgfr2 P253R* mutation on limb size but not on limb shape (Fig. 5a).

### *Fgfr2* Apert syndrome mutation leads to aberrant overexpression of *Dusp6* domains

We decided to obtain direct evidence of altered genetic regulation that could explain the observed limb phenotype by analyzing the shape dynamics of *Dusp6* expression, a direct target gene of *Fgf* signaling. As with limb shape, we first examined the gene expression of *Dusp6* in the embryos from the latest period. We found that at the “Late” period, *Dusp6* expression was already different between *Fgfr2*^*+/P253R*^ mutant mice and unaffected littermates. The differences were significant both in shape (Fig. 4e) and size (Fig. 5b and Table SI_3). In the limbs of unaffected mice, the *Dusp6* expression domain appeared as a thin domain underlying the apical ectodermal ridge, whereas in *Fgfr2*^*+/P253R*^ mutant mice the shape of the *Dusp6* domain was expanded in all directions (Fig. 4e). Accordingly, the volume of the *Dusp6* expression domain was significantly larger in Apert syndrome mice (Fig. 5b and Table SI_3), even when these mice presented significantly smaller limbs (Fig. 5a). The *Dusp6* expression domain thus grew disproportionately in the limbs of *Fgfr2*^*+/P253R*^ mutant mice in the latest period of development (Fig. 4e), which is consistent with a reported whole-body size reduction in *Fgfr2*^*+/P253R*^ Apert mice and the over-activation of *Fgfr2* signaling by the Apert syndrome mutation^27,44^.

At the “Mid late” period, an expanded *Dusp6* expression domain persisted on the dorsal and ventral sides of mutant limbs, but was reduced on the anterior and posterior sides (Fig. 4f). The overall volume of the *Dusp6* expression domains remained larger in *Fgfr2*^*+/P253R*^ mutant mice, but the difference was no longer statistically significant (Fig. 5b and Table SI_3).

At the “Mid early” period, the separation between unaffected and *Fgfr2*^*+/P253R*^ mutant mice was maintained in the PCA analysis (Fig. 4g). Unaffected mice showed a *Dusp6* expression domain expanded towards the anterior and posterior edges of the expression domain (Fig. 4g). In contrast, *Fgfr2*^*+/P253R*^ mutant mice did not show the extension and the posterior asymmetry of the *Dusp6* expression domain typical of normal limb development, suggesting a lack of differentiation in the *Dusp6* expression of mutant limbs (Fig. 4g).

Finally, the “Early” period was the only time point where we did not detect a significant separation between unaffected and *Fgfr2*^*+/P253R*^ mice (Fig. 4h). The PCA showed variation in the expression domains of *Dusp6*, with similar gene expression patterns in both shape (Fig. 4h) and size (Fig. 5b and Table SI_3) across all mice. Therefore, the first observation of an alteration in the gene expression pattern (Fig. 4g) occurred earlier than the alteration in the limb shape change (Fig. 4b). Our analyses provide evidence that differences in the *Dusp6* gene expression pattern occurred first, at the “Mid early period”, preceding the phenotypic limb differentiation, which occurred a few hours later in development, during the “Mid late period”. Overall, the time course showing the dynamics of limb and gene expression shape changes over development (Figs. 4, 5 and SI_3, 4 and 5) confirmed that the *Fgfr2* Apert syndrome mutation causes an aberrant overexpression of *Dusp6* early in development that could later lead to significant limb malformations.

### Altered *Dusp6* expression and limb dysmorphology are highly associated

Finally, we explored the correlation patterns between the limb phenotype and the gene expression pattern to further test whether altered *Fgf* signaling underlies the limb malformations induced by Apert syndrome *Fgfr2 P253R* mutation. First, we assessed the relationship between the size of the limbs and the volume of the *Dusp6* expression domain pooling all the forelimbs and hindlimbs and assessing the correlation between these two traits (Fig. 5c, d). The trend line showed that for the same limb size, *Fgfr2*^*+/P253R*^ mutant mice showed larger *Dusp6* expression domains, both in forelimbs (R^2^=0.4) and hindlimbs (R^2^=0.6). If the extension of the *Dusp6* expression only depended on limb growth, a high correlation between the size of the limb and the gene would be expected. However, the moderate correlation found here suggests that the size of the *Dusp6* gene expression is not dependent solely on limb size but is also influenced by other factors and could be under further genetic regulatory control.

Second, we assessed the morphological integration between the shape of the limbs and the shape of the *Dusp6* expression domain. The statistical analysis of the covariance pattern between these shapes can reflect the interaction of the phenotype and the gene expression pattern during limb development. As shown by analysis of additional genes expressed during limb development^15^, even when a gene is expressed within the limb, the shape of the limb and the shape of the gene expression domain are not correlated by definition, and the integration pattern can change from a strong association to no significant correlation within few hours of development^15^. The dynamics of the integration pattern can identify the key periods during which the expression of a gene is relevant for determining the shape of the limb. If the morphological integration is low, the expression of the gene will not be as relevant for the shape of the limb as if the integration is high. If the integration is low, the impact of the altered gene expression on the phenotype will be minimal. If the morphological integration is high, the impact of the genetic mutation will be maximized (i.e., changes of the gene expression pattern will produce changes in the limb shape). Our results showed that the shape of the limb and the shape of the *Dusp6* domain were indeed highly correlated (RV=0.88 in forelimbs; RV=0.91 in hindlimbs). This is evidence that altered *Fgf* signaling induced by the *Fgfr2 P253R* Apert syndrome mutation will have a direct effect on the limb phenotype.

By comparing the morphological integration pattern in mutant and unaffected littermates at different periods, we can test whether this interaction is maintained or disrupted by the disease during development. If the pattern or magnitude of morphological integration was different in mutant mice, it would reveal further mechanisms underlying the etiology of the disease. Our analyses showed that the pattern of morphological integration of the shape of the limbs and the shape of the *Dusp6* expression domains was similar in unaffected and *Fgfr2*^*+/P253R*^ mutant mice (Fig. 5e and Fig SI_6). Our results confirmed that the *Fgfr2 P253R* mutation does not disrupt the strong association between limb shape and the *Dusp6* expression domain. Therefore, the alteration of the *Dusp6* expression pattern caused by the *Fgfr2* mutation between E10 and E11.5 will produce the limb dysmorphologies associated with Apert syndrome. Overall, the high correlation between the shapes of the limb and the *Dusp6* expression domain provides further evidence that altered *Fgf* expression due to the *Fgfr2* mutation is strongly associated with limb defects in Apert syndrome.

## Discussion

By definition, to reveal the primary etiology of an abnormality requires going back in time to the earliest moment when abnormal development can be found. Typically, the earliest changes will be the most subtle, and so the most statistically sensitive techniques are necessary. The methods currently used to assess gene expression patterns are mainly qualitative and only focus on the shape and size differences that can be simply detected by eye. Therefore, slight changes in gene expression domains, even if they may have large effects on the phenotype^45^, can remain undetected. To reveal these subtle changes we have developed a precise method combining OPT and GM for quantifying embryo morphology and the underlying 3D gene expression patterns in a systematic, objective manner. This enables visualization and quantification of how the genotype translates into the phenotype during embryonic development, to compare normal and disease-altered patterns of genetic and phenotypic variation and, eventually, identify the origins of abnormal morphogenesis. This approach can further our understanding of the etiology of genetic diseases in research using animal models^46,47^, even in those that do not seem to recapitulate the human disease faithfully^48^.

Our study of the *Fgfr2 P253R* mouse model for Apert syndrome is an exemplary case of how quantitative assessment can overcome the shortcomings of traditional qualitative morphological assessment and lead to new discoveries. So far, the molecular and developmental mechanisms underlying the limb defects associated with Apert syndrome remained obscure, even when these limb abnormalities clinically differentiate Apert syndrome from other craniosynostosis syndromes (e.g. Pfeiffer, Crouzon and Saethre-Chotzen syndromes)^23,24,41^. Most Apert syndrome research focused on premature fusion of cranial sutures and craniofacial malformations^23,29–33^, clinical traits that are consistently phenocopied in mouse models. However, little research about the limb defects in Apert syndrome has been done using the same animal models, mainly because previous research reported the absence or subtle malformation of the limbs in the different mouse models for Apert syndrome^34–36^. Contrary to these previous results, our quantitative morphometric analyses demonstrate that the limbs of *Fgfr2*^*+/P253R*^ Apert syndrome mice present with significant defects that are detectable in newborn mice and can be traced back to early embryogenesis (Figs. 1, 4 and 5).

Our analyses provide insight into the genetic origins of these limb defects, showing that altered gene expression patterns in the *Fgf* signaling pathway precede and contribute to limb dysmorphogenesis in *Fgfr2*^*+/P253R*^ Apert syndrome mice. In fact, we detected that *Dusp6* expression patterns were different in unaffected and mutant littermates a few hours before the first limb dysmorphologies appeared (Fig. 4), and confirmed that limb shape and *Dusp6* expression patterns were highly correlated (Fig. 5e). The altered *Fgf* signaling observed was due to the *Fgfr2* P253R Apert syndrome mutation, which causes loss of ligand specificity of the *Fgfr2* isoform and increased affinity of various *Fgfs* to *Fgfr2*. Our analyses showed that over-activation of the *Fgf* signaling pathway results in more expanded (Fig. 4e-g) and larger (Fig. 5b) expression domains *of Dusp6*, a gene that acts as a negative-feedback control of *Fgf* signaling^49^. A delay in misregulation of the expression of *Dusp6* may explain the lack of antero-posterior asymmetry and the shape deficiencies observed in *Fgfr2*^*+/P253R*^ mutant mice (Fig. 4). We found evidence that limb shape and *Dusp6* expression were highly associated (Fig. 5c-e), but it is likely that other downstream genes of the *Fgf* signaling pathway also contribute to the limb shape malformations associated with Apert syndrome.

Our analyses also demonstrated that the embryonic limb defects persisted until birth (Fig. 1) and thus did not disappear over development. For instance, we found that *Fgfr2*^*+/P253R*^ Apert syndrome mice presented postnatal limb malformations involving the shape, length and volume of many bones of the forelimb, including the scapula, humerus, ulna, radius, metacarpals and phalanges (Fig. 1 and Table SI_1). Though subtle, these malformations suggest that further research into the origins and causes of limb dysmorphologies in Apert syndrome using these and other mouse models is warranted.

Our quantitative approach could be similarly applied to investigate other developmental defects and dysmorphologies ^50^. OPT and GM can be potentially used to analyze any organ and animal model that can be visualized during development using light microscopy, and any gene whose gene expression pattern can be detected by WISH and show a continuous expression domain over development^15^. We exemplified the method analyzing limb defects in mouse models, but it could be applied to malformations affecting other organs such as the face, the brain, the heart, etc, in mouse models as well as in any other vertebrate animal model, such as zebrafish, chicken and *Xenopus*. This is relevant as major developmental defects represent a leading cause of infant mortality and affect a small but relevant percentage of the population, severely compromising their quality of life^3^.

Through its increased quantitative sensitivity, our method has allowed us to reveal that the mouse model for Apert syndrome does indeed show early defects in limb development. We detected that dysregulation of an *Fgf* target gene precedes measurable changes in limb bud morphology, thus identifying a relevant component of its genetic etiology. Quantitative evaluation of size and shape should thus be performed before discarding any animal models as useful for investigating human congenital malformations^51^. Our method has the potential to become a high-throughput tool for biomedical research, providing insight into the processes that cause malformations and lead to malfunction, which is essential for understanding diseases and discovering potential therapies.

## Methods

### Mouse model

We analysed the *Fgfr2*^*+/P253R*^ Apert syndrome mouse model, an inbred model backcrossed on C57BL/6J genetic background for more than 10 generations carrying a mutation that in humans with Apert syndrome is associated with more severe syndactyly. This gain of function mutation, which is embryonically lethal in homozygosis, involves a proline to arginine amino acid change at position 253 of the *Fgfr2* protein that alters the ligand binding specificity of the receptor and causes a stronger receptor signaling. Further details of the mouse model on generation of targeting construct can be found elsewhere^34^. All the experiments were performed in compliance with the animal welfare guidelines approved by the Pennsylvania State University Animal Care and Use Committees (IACUC46558, IBC46590).

### Micro-CT imaging

High resolution micro-computed tomography (μCT) scans were acquired by the Center for Quantitative Imaging at the Pennsylvania State University (www.cqi.psu.edu) using the HD-OMNI-X high-resolution X-ray computed tomography system (Varian Medical Systems, Palo Alto, CA). Pixel sizes ranged from 0.01487 to 0.01503 mm, and all slice thicknesses were 0.016 mm. Image data were reconstructed on a 1024×1024 pixel grid as a 16-bit TIFF. To reconstruct forelimb morphology from the μCT images, isosurfaces were produced with median image filter using the software package Avizo 6.3 (Visualization Sciences Group, VSG) (Fig. 1a-f).

### Morphometrics in P0 mice

We assessed forelimb morphology at P0 using unaffected (n=10) and mutant (n=12) littermates of Apert syndrome mouse models. A set of 16 landmarks were collected on each left forelimb, including points on the main bones of the forelimb (phalanges, metacarpals, radius, ulna, humerus, scapula and clavicle), as shown in Fig. 1b-f. To minimize measurement error, each landmark was collected twice by the same observer restricting the deviations between the two trials to 0.05 mm.

At P0, we estimated the dimensions of the long bones of the forelimbs using the 3D coordinates of the landmarks located at the proximal and distal ends of the bones (Fig. 1b-f). We also estimated the bone volumes from volume data collected from the μCT scans. To assess size differences in bone length and bone volume between mutant and unaffected P0 mice of the *Fgfr2*^*+/P253R*^ Apert syndrome mouse model, we performed a two-tailed one-way ANOVA on those variables showing a normal distribution, and the non-parametric Mann-Whitney U-Test on those variables that deviated from a normal distribution. The shape of the humerus and the scapula was comparatively assessed in unaffected and *Fgfr2*^*+/P253R*^ Apert syndrome littermates using Geometric Morphometrics. Shape information was extracted using a General Procrustes Analysis (GPA)^52^, in which configurations of landmarks are superimposed by shifting them to a common position, rotating and scaling them to a standard size until a best fit of corresponding landmarks is achieved^53^. The resulting Procrustes coordinates from the GPA were the input for further statistical analysis to compare the shape of the bones in unaffected and *Fgfr2*^*+/P253R*^ mice. A Principal Component Analysis (PCA) was used to explore the morphological variation within each bone. PCA performs an orthogonal decomposition of the data and transforms variance covariance matrices into a smaller number of uncorrelated variables called Principal Components (PCs), which successively account for the largest amount of variation in the data^20^. Each specimen is scored for every principal component and the specimens can be plotted using these scores along the morphospace defined by the principal axes.

### WISH and OPT scanning

To examine early embryonic mouse limb development in Apert syndrome we bred 4 litters of the *Fgfr2*^*+/P253R*^ Apert syndrome mouse model and collected them between E10.5 and E11.5. In total, 32 mouse embryos were harvested and classified by PCR genotyping into unaffected (n=16) and mutant (n=16) littermates (see Fig. SI_1 and Table SI_2 for further details on sample size and composition). *Dusp6* gene expression was assessed by Whole-Mount-In-Situ Hybridization (WISH). Mouse embryos were dissected in cold phosphate-buffered saline, 0.1% tween 20 (PBT), fixed overnight in 4% paraformaldehyde (Sigma), dehydrated in a graded PBT/methanol series and stored at −20ºC in methanol. The mouse embryos recovered their original size after rehydration in decreasing series of methanol/PBT. WISH was carried out using *Dusp6* antisense RNA probes labelled with digoxigenin-UTP (Roche), following standard protocols^7^. Alkaline phosphatase coupled anti-digoxigenin (anti-DIG-AP, Roche) and NBT/BCIP staining (Roche) were used to reveal the expression pattern for *Dusp6*. To minimize variations during experimental procedures, all embryos were processed systematically within the same batch, processing unaffected and mutant littermates from different litters in separate tubes, but simultaneously using the same probe, timings and concentration reagents.

After embedding in agarose, dehydrating in methanol and chemically clearing the samples with benzyl alcohol and benzyl benzoate (BABB), the stained whole embryos were scanned with both fluorescence and transmission light with a CFP (Cyan Fluorescent Protein) filter using our home-build OPT imaging system mounted on a Leica MZ 16 FA microscope^5^. The embryos were 3D-reconstructed from the resulting 2D images using Matlab and visualized using Amira 6.3 (Visualization Sciences Group, FEI). From the OPT fluorescence scans we produced 3D reconstructions of the embryo surface and we dissected the available right and left fore- and hind limbs of each specimen, resulting in a sample of 101 embryonic limbs (Table SI_2). From the OPT transmission scans, we recovered the *Dusp6* expression domain. As *Dusp6* is expressed in a fuzzy spatial gradient, as already shown by other genes^15^, we used 3D multiple thresholding to visualize the gene expression domain at different intensities (Fig. 3 and Video SI_1). To comparatively analyse the gene expression domains across the sample, we inspected for each limb the whole range of threshold values under which the gene expression could be visualized, from the threshold showing its first appearance to the threshold under which it disappeared and was no longer detectable. We analysed the 3D reconstruction based on a threshold computed as 1/3 of the last grey value showing the *Dusp6* expression domain, which displayed a *Dusp6* domain at high gene expression (Fig. 3, step 3). Finally, we obtained 80 limbs (46 forelimbs and 34 hindlimbs) with associated gene expression patterns for *Dusp6.*

### Embryo staging

To account for breeding and developmental variation, individual limb buds were staged using our publicly available web-based staging system (http://limbstaging.crg.es)^43^. Considering the spline curve along the outline of the limb, this tool provides a stage estimate with a reproducibility of ± 2 hours. According to the staging results, the different mouse litters were ordered following a continuous temporal sequence from E10 to E11.5 (Fig. SI_1). To minimize high developmental variation within and among litters of mice, hindlimbs and forelimbs from each litter were analysed separately, except for those from two litters from the earliest stage that were pooled into the same group because their temporal distribution completely overlapped, as shown in Fig. SI_1.

### Morphometrics from E10 to E11.5

To capture the size and shape of the limbs and the expression domains of *Dusp6*, we collected a set of anatomical landmarks as well as curve and surface semilandmarks (Fig. SI_2), as recommended in structures devoid of homologous landmarks. Semilandmarks are mathematical points located along a curve^54^ or a surface^55^ within the same object that can be slid to corresponding equally spaced locations across the sample. We used Amira 6.3 (Visualization Sciences Group, FEI) to record the anatomical landmarks and Viewbox 4 (dHAL software, Kifissia, Greece) to construct a limb template of surface and curve semilandmarks and to interpolate them onto each target shape (Fig. SI_2).

The 3D landmark coordinates defining the shape of the limb and the *Dusp6* expression domain were analysed using Procrustes-based landmark analysis^54^. Semilandmarks were allowed to slide in the GPA by minimizing the bending energy^54–56^. Quantitative shape analyses based on PCA were performed as explained above.

We estimated the size of the limb as the centroid size, calculated as the square root of the summed squared distances between each landmark coordinate and the centroid of the limb configuration of landmarks^53^. The volumes of the *Dusp6* domains were estimated from the 3D reconstructions of the OPT scans. Size differences in limb size and gene volume between mutant and unaffected embryonic mice were tested for statistical significance using a two-tailed Welch Two Sample t-test.

We quantified the integration between the limb and the *Dusp6* expression pattern and produced visualizations of the patterns of associated shape changes between them using Two-Block Partial Least Squares analysis (PLS)^57^. This method performs a singular value decomposition of the covariance matrix between the two blocks of shape data (i.e., the limb and the *Dusp6* expression domain). Uncorrelated pairs of new axes are derived as linear combinations of the original variables, with the first pair accounting for the largest amount of inter-block covariation, the second pair for the next largest amount and so on. The amount of covariation is measured by the RV coefficient, which is a multivariate analogue of the squared correlation^58^. Statistical significance was tested using permutation tests under the null hypothesis of complete independence between the two blocks of variables. Separate analyses for each developmental period, as well as for forelimbs and hindlimbs of all stages were computed.

All the analyses were performed using R (R Development Core Team, 2013; http://www.R-project.org); the R package geomorph^59^ (http://cran.r-project.org/web/packages/geomorph), SPSS Statistics 22 (IMB, 2013) and MorphoJ^60^.

## Acknowledgements

We acknowledge support of the Spanish Ministry of Economy and Competitiveness to the EMBL partnership* and ‘Centro de Excelencia Severo Ochoa, and the CERCA Programme / Generalitat de Catalunya. We acknowledge the support of the CERCA Programme / Generalitat de Catalunya. We acknowledge Ethylin Wang Jabs for access to the *Fgfr2*^*+/P253R*^ Apert syndrome mouse model. The research leading to these results received funding from the following grants: European Union Seventh Framework Program (FP7/2007-2013) under grant agreement Marie Curie Fellowship FP7-PEOPLE-2012- IIF 327382, National Institutes of Health NICHD P01HD078233 and NIDCR R01DE02298, and a Burroughs-Welcome Fund 2013 Collaborative Research Travel Grant.

## Author contributions

NM-A, JS and JR conceived the project. JR contributed with experimental mouse models; AR-M, JS, LS and KK conducted experiments and scanning; and NM-A, RM, JS, SMP and MY collected and analysed the data. NM-A and JS wrote the paper and all authors critically reviewed the manuscript.

